# Confounding factors in algal phosphorus limitation experiments

**DOI:** 10.1101/298281

**Authors:** Whitney S. Beck, Ed K. Hall

## Abstract

Assessing algal nutrient limitation is critical for understanding the interaction of primary production and nutrient cycling in streams, and nutrient diffusing substrate (NDS) experiments are often used to determine limiting nutrients such as nitrogen (N) and phosphorus (P). Unexpectedly, many experiments have also shown decreased algal biomass on NDS P treatments compared to controls. To address whether inhibition of algal growth results from direct P toxicity, NDS preparation artifacts, or environmental covariates, we first quantified the frequency of nutrient inhibition in published experiments. We also conducted a meta-analysis to determine whether heterotrophic microbial competition or selective grazing could explain decreases in algal biomass with P additions. We then deployed field experiments to determine whether P-inhibition of algal growth could be explained by P toxicity, differences in phosphate cation (K vs. Na), differences in phosphate form (monobasic vs. dibasic), or production of H_2_O_2_ during NDS preparation. We found significant inhibition of algal growth in 12.9% of published NDS P experiments as compared to 4.7% and 3.6% of N and NP experiments. The meta-analysis did not show enhanced heterotrophy on NDS P treatments or selective grazing of P-rich algae. Our field experiments did not show inhibition of autotrophic growth with P additions, but we found significantly lower gross primary productivity (GPP) and biomass-specific GPP of benthic algae on monobasic phosphate salts as compared to dibasic phosphate salts, likely because of reduced pH levels. Additionally, we note that past field experiments and meta-analyses support the plausibility of direct P toxicity or phosphate form (monobasic vs. dibasic) leading to inhibition of algal growth, particularly when other resources such as N or light are limiting. Given that multiple mechanisms may be acting simultaneously, we recommend practical, cost-effective steps to minimize the potential for P-inhibition of algal growth as an artifact of NDS experimental design.

## Introduction

Benthic algal production provides an important energy source to higher trophic levels [1], and in low productivity streams, growth of macroinvertebrate and fish grazers may be limited by the availability of algal food resources [2]. Freshwater algal growth is often limited by the availability of nitrogen (N), phosphorus (P), or both nutrients [3, 4], but human activities are increasing N and P inputs to streams via sources such as wastewater treatment effluent, agricultural runoff, and atmospheric N deposition [5]. These excess nutrients may result in harmful levels of algal biomass that degrade ecological habitat [5], stream aesthetics [6], and drinking water quality [5]. Identifying nutrients that limit algal productivity in individual stream reaches can inform stream management plans that promote human and ecosystem health.

For over thirty years, nutrient diffusing substrate (NDS) experiments have been used to determine nutrient limitation of benthic algal communities [7, 8]. Nutrient diffusing substrate experiments are constructed by filling a small vessel (e.g., a plastic vial or clay pot) with agar and a nutrient solute of choice (e.g., KH2PO4), and comparing growth with a paired control vessel containing only agar [9]. Differences in algal responses can then be compared across the NDS treatments. Contrary to the long-held paradigm that P frequently limits algal productivity in freshwater ecosystems [10], one of the first published NDS experiments observed that treatments with 0.5 M P had lower algal biomass than treatments with 0.05 M P [7]. Many studies have since found that algal biomass can be inhibited by P as compared to controls (e.g., [11, 12]), and it is unclear how often or why this phenomenon occurs. Whereas P may not always enhance growth due to limitation by N or other resources (light, Fe, etc.), it is surprising that increasing levels of P in NDS can result in decreased algal biomass relative to the control treatment. These results suggest that artifacts associated with NDS experiments may be leading to the underreporting or misrepresentation of P-limitation in freshwater ecosystems.

Several hypotheses have been introduced to explain why addition of P in NDS would result in a decrease of algal biomass. As a macronutrient, P is required for algal growth and maintenance [13], but high P concentrations may result in direct physiological toxicity and this has been hypothesized as a reason for observed P-inhibition [7]. While mechanisms for this toxicity in algae are unclear, excessive P concentrations in growth media of terrestrial plants can negatively affect the availability, uptake, and metabolic processing [14] of Fe [15], K [15], and Z [16], leading to deficiency of these essential nutrients and slowing or inhibiting plant growth. Other hypothesized mechanisms for P-inhibition of algal growth have focused on artifacts related to preparation of the NDS, including: 1) phosphate cation type (K vs. Na) and toxicity, 2) phosphate form (monobasic vs. dibasic), and 3) H_2_O_2_ production from phosphate reacting with agar during autoclaving. A limited number of laboratory studies have tested whether high concentrations of the phosphate salt cations (K^+^, Na^+^, H^+^) may inhibit algal growth. These studies have shown that K phosphates and KCl are toxic to algae at lower concentrations than Na phosphates and NaCl [17, 18]. The phosphate form may also influence algae, as monobasic forms (KH_2_PO_4_ and NaH_2_PO_4_) tend to have lower NDS effect sizes than dibasic forms (K_2_HPO_4_ and Na2HPO4 [19]). This suggests algae are either inhibited by acidic pH levels (induced by monobasic forms) or are experiencing cation limitation (alleviated by dibasic forms). Lower pH may influence algae directly by changing concentrations of H^+^ around the cell or indirectly through the effects of pH on metal toxicity or nutrient availability (e.g., via slowed nitrification rates or binding of P by Al [20]). Finally, the preparation of the NDS media may affect how the P treatments influence algal growth. Autoclaving phosphate and agar together produces H_2_O_2_, a toxin that may inhibit microbial growth [21]. It has been suggested that the common NDS construction method of combining the two compounds on a hotplate could also produce the same result [9] thus leading to inhibition of algal growth in treatments that contain P; however, to our knowledge this has never been directly tested.

Beyond these direct artifacts of NDS preparation, there may also be a series of indirect effects of adding phosphate to NDS that could inhibit algal growth. For example, P could disproportionately stimulate heterotrophic microbes [22] and increase competition between autotrophs and heterotrophs for other limiting nutrients, ultimately suppressing algal growth. It is also possible that P amendments may induce additional top-down pressure if insect grazers selectively graze P-rich algal biofilms [23], resulting in lower algal biomass on NDS P treatments as compared to controls [11]. Selective foraging has been supported by a theoretical analysis [24], and laboratory experiments provide some additional support that grazers can engage in P-specific foraging [23, 25].

Determining why P-inhibition of algal growth has been observed in NDS experiments is not only an important methodological question but one with important implications for an ecological understanding of lotic ecosystems. Each of the previously-described mechanisms may potentially affect the response of algae to P treatments in NDS experiments, but there is no single study that simultaneously evaluates each mechanism. We used quantitative analyses of published data and our own field experiments to investigate how frequently and why NDS P treatments inhibit algal growth. First, we surveyed the literature to determine how frequently significant inhibition of algal growth was reported for P, N, and NP treatments. We also completed a meta-analysis to determine if there was consistent evidence for heterotrophic microbial competition or top-down grazing control that could be leading to P-inhibition of algal growth across multiple study systems. Finally, we completed field experiments to directly address the effects of NDS preparation on several common response variables used to evaluate algal growth: chlorophyll *a* (a measure of algal biomass), ash-free dry mass (AFDM, a measure of total biofilm organic matter), a calculated index of autotrophy (AI), gross primary productivity (GPP), and biomass-specific GPP (GPP/chlorophyll a).

## Methods

### Quantitative Review

To explicitly quantify how often nutrient treatments inhibit algal growth in NDS experiments, we used the database assembled by Beck et al. [19]. Briefly, this database includes 649 NDS experiments from 1985-2015 that used algal biomass (chlorophyll *a*) as a response variable. The database was previously used to determine overall effect sizes of P, N and NP additions and to quantify the influence of over thirty experimental, environmental, and geographic covariates. However, in this study our goal was to determine how many individual experiments detected significantly lower algal biomass levels on P treatments (n=534), N treatments (n=553), and NP treatments (n=591) as compared to controls (α=0.05). Although we could have used a meta-analysis approach, we used separate two-tailed t-tests for each NDS experiment to better represented how investigators analyzed data in individual studies.

To determine whether P treatments significantly influenced the proportion of autotrophy in microbial communities and whether grazers selected for algal biomass on NDS P treatments, we used meta-analysis models (see below). To assess whether P additions changed algal-heterotrophic interactions, we identified any experiments from the previously compiled database [19] that also reported AFDM as a response variable, as a proxy for total biofilm organic matter. For this study, we extracted AFDM data using Webplot Digitizer version 3.12 [26], and calculated an autotrophic index (AI) for each experiment and treatment as follows:

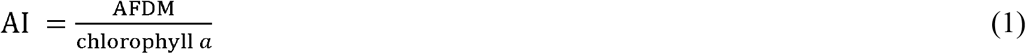

We interpret lower values of the AI as a higher proportion of autotrophy in the microbial community [22]. For NDS P treatments we calculated AI log response ratios (LRRs, a measure of effect size) as:

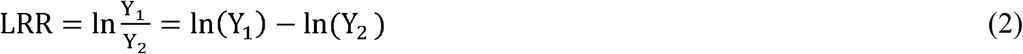

Where Y_1_ is the mean AI from the P treatments and Y_2_ is the mean AI from the control treatments in a given experiment [27]. LRRs greater than zero indicate there is a positive P treatment effect, but LRRs less than zero indicate an inhibitory effect of the P treatment as compared to controls. We also calculated the variance of effect sizes using:

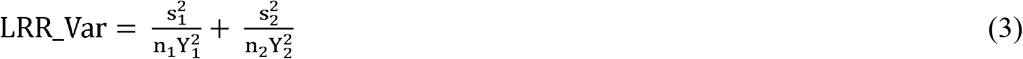

Where s_1_ and s_2_ are the standard deviations of the P and control treatments, and n_1_ and n_2_ are the number of P and control replicates [27]. To determine whether P additions influence grazer selection of algal biofilms, we identified NDS experiments that also incorporated grazer exclusion treatments. We calculated LRR and LRR_Var metrics for P treatments compared to controls in grazed and ungrazed plots (Equations 2-3).

We used the metafor package in R to build meta-analysis models based on experimental effect sizes and variances [28]. Meta-analysis models account for variability within and among experiments by weighting effect sizes by their variances [28]. In all models, we used “site” as a random effect, to account for the correlated effects of experiments deployed in the same stream reach [29]. Models were used to determine how P additions influenced AI effect size, and to determine how grazers influenced P treatment effects on algal biomass. For the grazer models, we included grazer treatment as a covariate in the meta-analysis models [28].

## Experiments

To investigate how NDS preparation methods influence P-inhibition of algal growth, we prepared and deployed a series of complementary NDS experiments in a sub-alpine stream. To address the influence of cation type, monobasic vs. dibasic phosphate form, and the potential for H_2_O_2_ formation, the first two experiments involved crossing four different phosphate chemicals (KH_2_PO_4_, K_2_HPO_4_, NaH_2_PO_4_, or Na_2_HPO_4_ all at 0.1 M concentrations) with two laboratory heating methods (boiling agar and phosphate together vs. separately) for a total of eight preparation treatments (Fig 1). In a third experiment, we again crossed the four phosphate compounds with two laboratory heating methods but to assess direct toxicity of excess P, we also used two different concentrations of P (0.05 M and 0.5 M) for a total of 16 preparation treatments. Concentrations were chosen based on the most commonly used NDS concentrations. Furthermore, Fairchild et al. [7] previously observed a difference in algal biomass responses to 0.05 M and 0.5 M of P.

**Fig 1.**
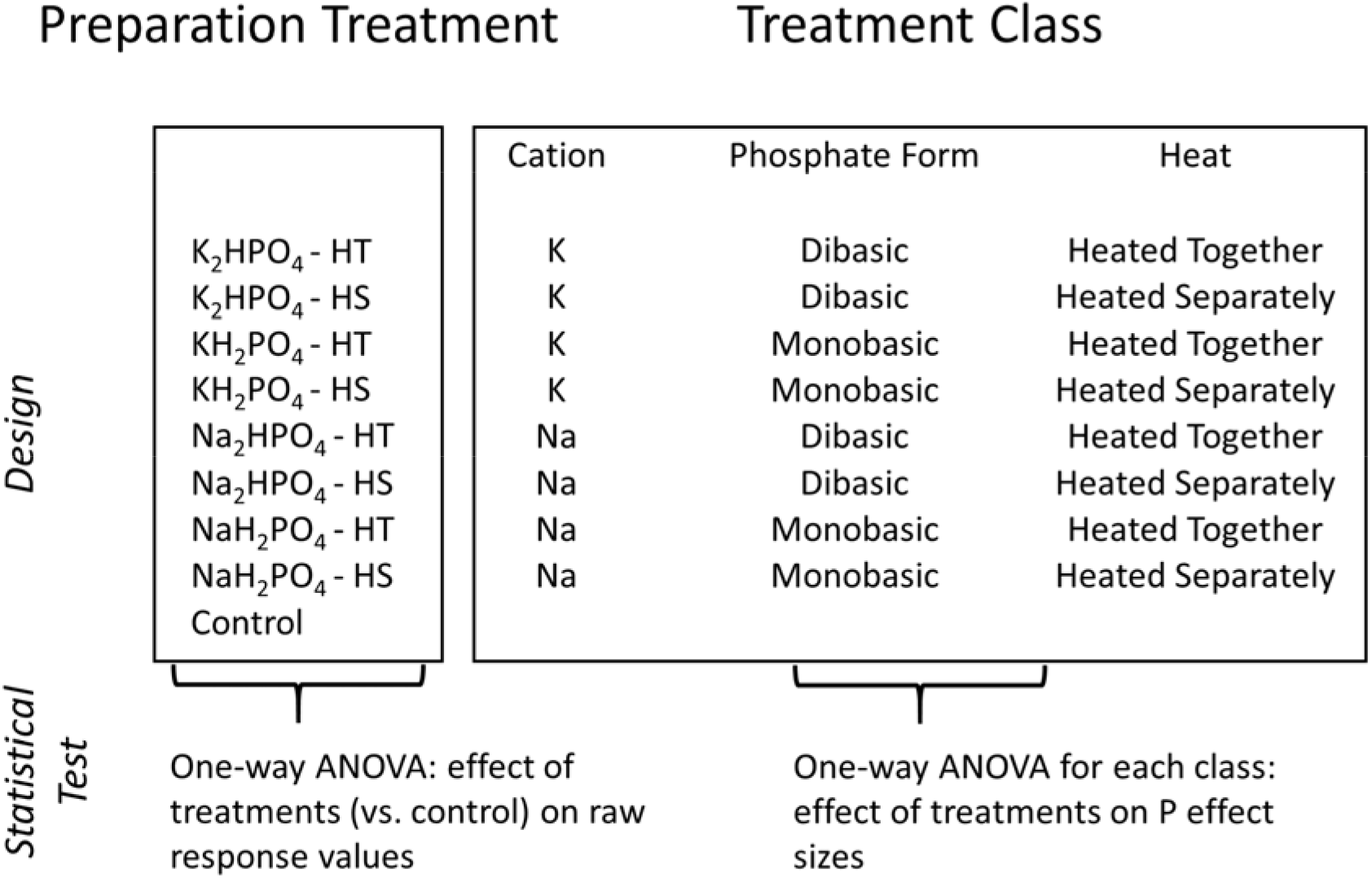
Field experiment preparation methods and treatment classes. Four different phosphate chemicals crossed with laboratory heating methodology were deployed in two NDS experiments in 2016, for a total of eight preparation treatments. The eight treatments were grouped by three treatment classes including cation, phosphate form, and heating method. The same preparation treatments were crossed with two different phosphate concentrations in a 2017 experiment, for a total of sixteen preparation treatments. The sixteen treatments were grouped by four treatment classes including cation, phosphate form, heating method, and concentration. Heated Together = phosphate and agar boiled together, Heated Separately = agar boiled and phosphate added at pouring temperature of 55-65° C.

We constructed NDS according to standard methods in the literature [9]. Briefly, we boiled 2% agar with deionized water, poured the solutions into 30 mL vials (Item #66159, U.S. Plastic Corps, Lima OH), and capped the cooled agar with fritted glass discs (5.7 cm^2^, Item #C4505, EA Consumables, Pennsauken, NJ). A plain agar solution was used for the control, and the specified phosphate salt was added to the agar according to the experimental design described above (Fig 1). For the “heated together” treatment, we boiled the agar and the phosphate salt together. For the “heated separately” treatment, we added the phosphate salt once the agar had cooled to handling temperature (55-65° C) and mixed it thoroughly with a magnetic stir plate before solidification.

While preparing the NDS treatments we also measured pH, a putative mechanism for how phosphate form mediates P-inhibition of algal growth. We tested the effects of phosphate compound and heat treatment on agar and water pH in the lab. To do this we used litmus pH test strips to measure the pH of cooling agar for four vials from each phosphate treatment and the control. To measure effects of phosphate compound and heat treatment on water pH, we constructed two replicate NDS representing the four phosphate compounds crossed with two heat treatments, as well as a control. These NDS were placed in separate plastic Ziploc^®^ bags with 1 L of distilled water. We used a multimeter with a pH probe (Thermo Fisher Scientific Orion Star™ A329, Waltham, MA) to measure the initial pH and remeasured the pH after 24 hours.

We deployed all NDS experiments at Little Beaver Creek, a low-order, open-canopy stream in the mountains of the Roosevelt National Forest in Colorado (40.625° N, −105.527° W). Experiments 1 and 2 were deployed in summer 2016, while experiment 3 was deployed in summer 2017 (S1 Table). Previous NDS experiments in summer 2015 showed that Little Beaver was co-limited by N and P, but primarily N-limited, with P treatments causing inhibition of algal biomass relative to the control (Beck, unpublished). We randomized 6 replicates of each treatment and attached 6-8 individual NDS vials to plastic L-shaped bars (Item #45031, U.S. Plastic Corp, Lima, OH) that were anchored into the streambed using metal stakes. Upon deployment, collection, or both, we measured in-stream pH, conductivity, and temperature using a multimeter and probes (Thermo Fisher Scientific Orion Star™ A329, Waltham, MA); collected duplicate filtered (0.45 μm Type A/E filters, Pall Corporation, Port Washington, NY) water samples in 60 mL nalgene™ bottles for nutrient analysis; measured canopy cover using a densiometer (Forestry Suppliers, Jackson, MS); and measured flow using a Marsh McBirney meter (Hach, Loveland, CO). We used a MiniWater^®^ 20 meter (Schiltknecht, Switzerland) to measure 2.5 cm-scale current velocity above three evenly-spaced vials on each L-bar. We measured NO_3_^-^ using the Cd reduction method [30] and orthophosphate using the ascorbic acid method [31] on an ALPKEM^®^ Flow Solution IV autoanalyzer (O.I. Analytical, College Station, TX).

We analyzed primary production rates on NDS discs after each in-stream experiment [9]. Briefly, upon collecting the discs, we immediately placed them in 50 mL centrifuge tubes with unfiltered river water, capping the tubes underwater to exclude any air bubbles. Tubes were stored on ice during transport back to the laboratory (less than two hours), where we used filtered (0.45 μm Type A/E filters, Pall Corporation, Port Washington, NY) river water to run light and dark incubations with the NDS discs. We measured the initial temperature and dissolved oxygen (DO) values of the water before light and dark incubations using a ProDO meter (YSI, Yellow Springs, OH). We also included four “blank” tubes with water only, to control for background changes in DO because of temperature changes or exposure to atmospheric oxygen during the measurement process. During light treatments, tubes were incubated in sunlight for two hours in the afternoon, then we measured the ending DO concentration in each tube and ending temperature in several representative tubes. For dark incubations, we replaced the filtered water and incubated tubes in the dark for two hours, after which we measured the final DO concentrations in each tube and ending temperature in several representative tubes.

We calculated net primary productivity (NPP) as the increase in oxygen during the light incubations, correcting for the change in the blank stream water tubes. We calculated respiration as the decrease in oxygen during the dark incubations, correcting for the change in the blank stream water tubes. Gross primary productivity was calculated as follows, then all variables were standardized by disc area and time:

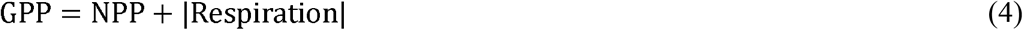

After the incubations, we immediately placed the discs in black film canisters and extracted chlorophyll *a* for 12-24 hours using buffered 90% ethanol. We measured chlorophyll *a* with an acidification correction [32] using an Aquafluor^®^ fluorometer (Turner Designs, San Jose, CA). We calculated biomass-specific GPP (GPP/ chlorophyll *a)* as an additional response metric for each disc.

For experiments 1 and 2, we saved all liquid slurry from the chlorophyll *a* extractions and filtered the liquid through a pre-combusted filter (500° C for one hour, 0.45 μm Type A/E, Pall Corporation, Port Washington, NY). We then used both the filter and glass disc associated with each NDS to measure AFDM [22]. For experiment 3, we allowed the chlorophyll *a* extraction slurry to evaporate in a weigh boat under a fume hood and used all remaining material and the glass disc to measure AFDM. We dried the samples for 48 hours at 50° C, pre-weighed their masses, and combusted them at 500° C for one hour in a muffle furnace. We then rehydrated the discs with deionized water, dried them for another 48 hours at 50° C, and weighed the final masses. This procedure was to account for any water that might have been lost from clay particles in the muffle furnace. The difference in weights was calculated as AFDM. We also used the chlorophyll *a* and AFDM measurements to calculate AI as described in Equation 1.

## Statistical Analyses for Experiments

We completed all statistical analyses in R version 3.5.0 (R Core Team 2018). To test the effect of treatment classes on the pH of the NDS agar and water incubated with the NDS, we used two-way ANOVAs with phosphate form (monobasic vs. dibasic) and heating method as factors. We compared means using post-hoc Tukey’s HSD tests (α=0.05).

Because the field experiments comprised an incomplete factorial design (i.e., crossed treatments and a separate control, Fig 1), we used three separate approaches to analyze the results. First, to determine whether P preparation treatments significantly stimulated or inhibited response variables relative to controls, we used one-way ANOVAs with post-hoc Dunnett tests to compare each of the eight P preparation treatments (Fig 1) with the controls. The Dunnett test is a multiple comparison procedure that compares individual treatment means to control means while maintaining a family-wise error rate that is below *α* (α = 0.05 in this study). We used only data from experiments 1 and 2 and included experiment as a fixed effect block to control for experiment-specific artifacts. Second, to determine whether the treatment classes (cation type, phosphate form, and heat, Fig 1) influenced NDS P effect sizes, we used data from experiments 1 and 2 to run separate one-way ANOVAs for each treatment class and response variable (algal biomass, AFDM, AI, GPP, and biomass-specific GPP). We included experiment as a fixed effect block, and the model response variables were P treatment values of each response variable divided by their respective experimental control means [9]. Third, to determine whether treatment classes interacted with P concentration to influence NDS P effect sizes, we used data from experiment 3 to run separate two-way ANOVAs crossing each treatment class with concentration.

## Results

### Quantitative Review

In our analysis of 649 of experiments from the literature we found that algal biomass was more commonly inhibited by NDS P treatments than either N or NP treatments. Phosphorus additions produced a significant negative effect in 12.9% of experiments. However, N and NP additions produced a significant negative effect in only 4.7% and 3.6% of experiments, respectively (both within the commonly assumed type I error rate of 5%).

We next looked for published evidence of hypothesized biological mechanisms that would explain inhibition of algal growth by P. To address the potential for heterotrophic suppression of autotrophs in microbial communities on NDS P treatments, we identified 45 experiments from 11 studies where an AI effect size could be calculated (S1 References). However, the meta-analysis of AI effect sizes showed neither a positive nor negative response to P treatments (Fig 2). We found even fewer examples of studies that could be used to examine nutrient additions in conjunction with grazer exclusions. Only five experiments from three studies reported the effects of grazer exclusions on P treatments in NDS experiments (S2 References). In these studies, algal biomass effect sizes on the P treatments were not significantly influenced by the presence or absence of grazers (Fig 2, Q_M_ = 2.594, p=0.107).

**Fig 2.**
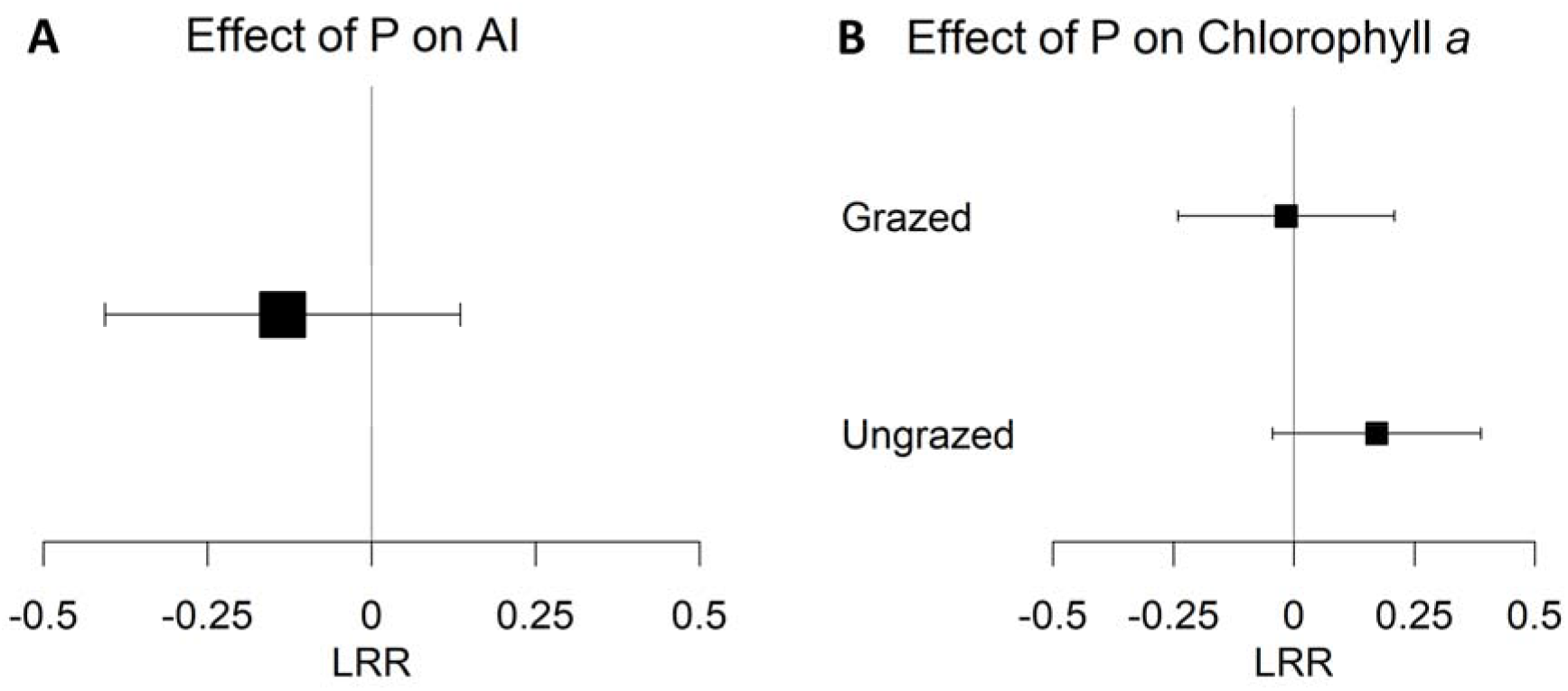
Meta-analysis model results. (A) Results of a meta-analysis model testing the significance of NDS P treatment autotrophic index (AI) effect sizes (n=45 experiments), where lower values indicated a higher proportion of autotrophy in microbial communities. (B) Results of a meta-analysis model testing the effect of grazing on NDS P treatment algal biomass (chlorophyll *a*) effect sizes (n=5 experiments per grazing treatment). Squares are log response ratio (LRR, see equation 2) mean estimates surrounded by 95% confidence interval bars.

**Fig 3.**
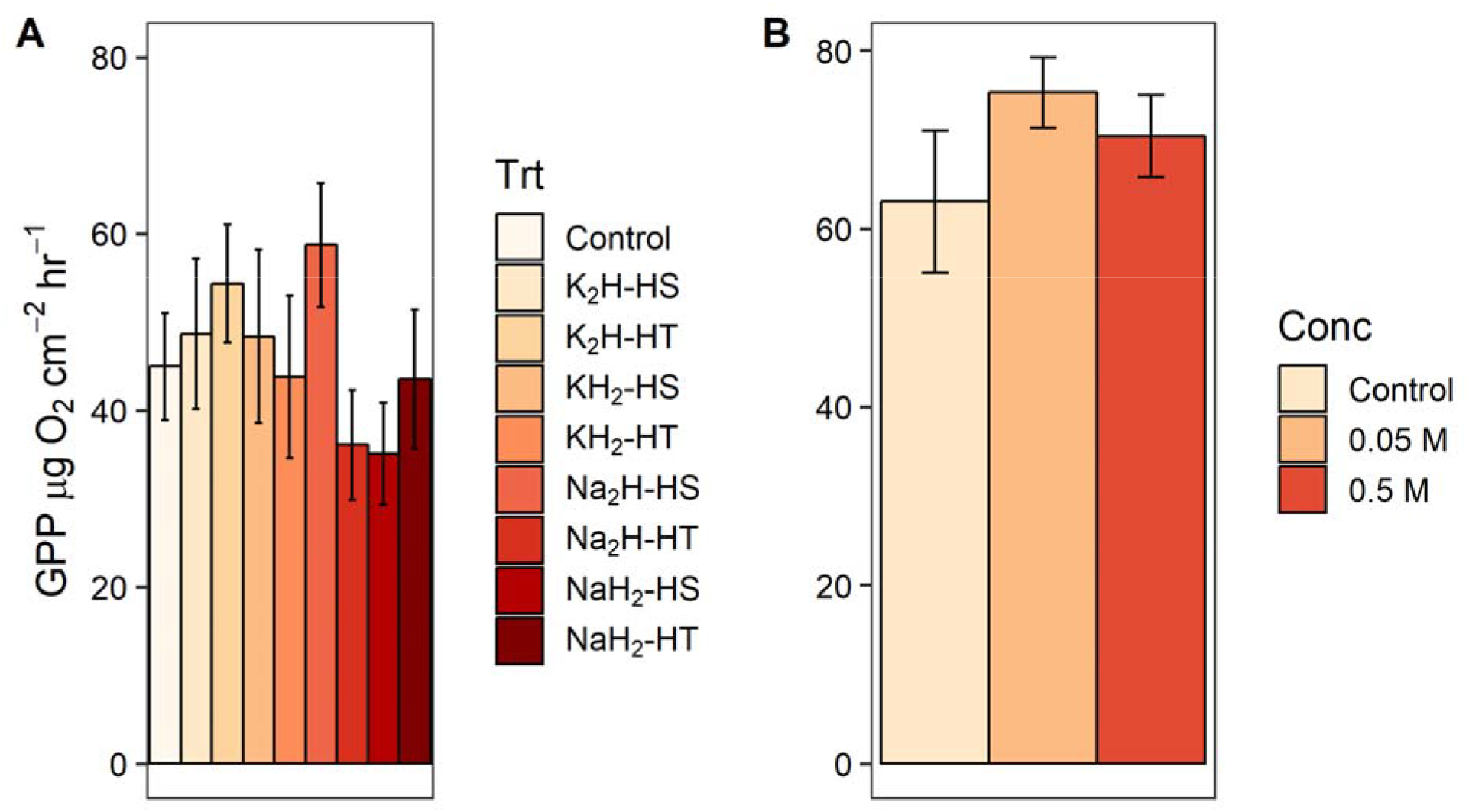
Effect of experimental preparation treatments on gross primary productivity (GPP). Gross primary productivity means ± 1 standard error from nutrient diffusing substrate (NDS) field experiments. (A) Results from experiments 1-2 (total n=103), where treatments consisted of crossing four chemicals (K_2_HPO_4_, KH_2_PO_4_, Na_2_HPO_4_, and NaH_2_PO_4_) and two heat treatments (agar and phosphate heated together vs. heated separately, denoted as “HT” and “HS”). An agar-only control treatment was included in each experiment. (B) Results from experiment 3 (total n=104), where treatments included the same factors as experiments 1-2, except low (0.05 M) and high (0.5 M) phosphate concentrations were included as an additional factor. Gross primary productivity values by concentration are presented by averaging over all other factors.

**Fig 4.**
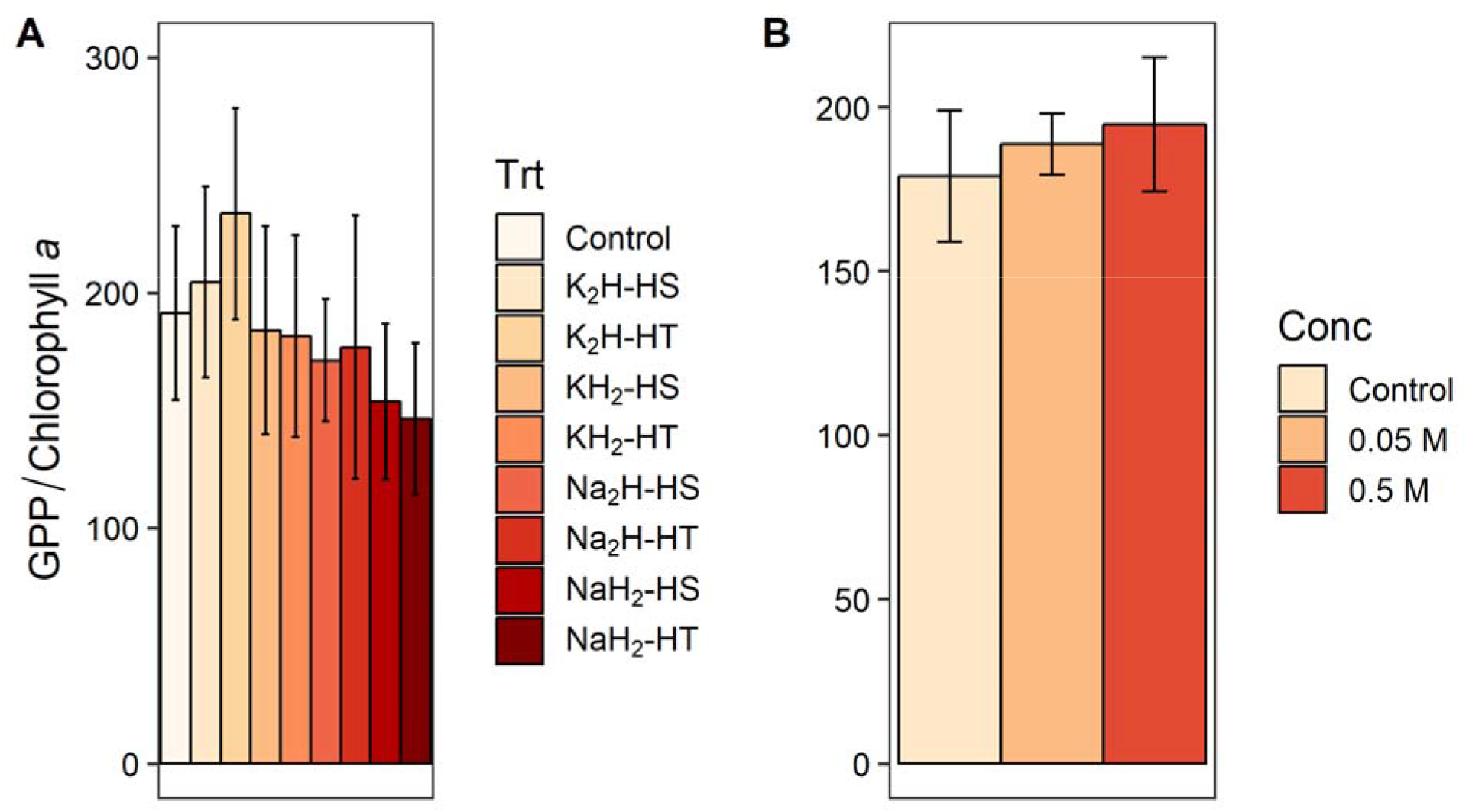
Effect of experimental preparation treatments on biomass-specific gross primary productivity (GPP). Biomass-specific GPP means ± 1 standard error from NDS field experiments (see Figure 2 caption). (A) For experiments 1-2, total n=102. (B) For experiment 3, total n=103.

**Fig 5.**
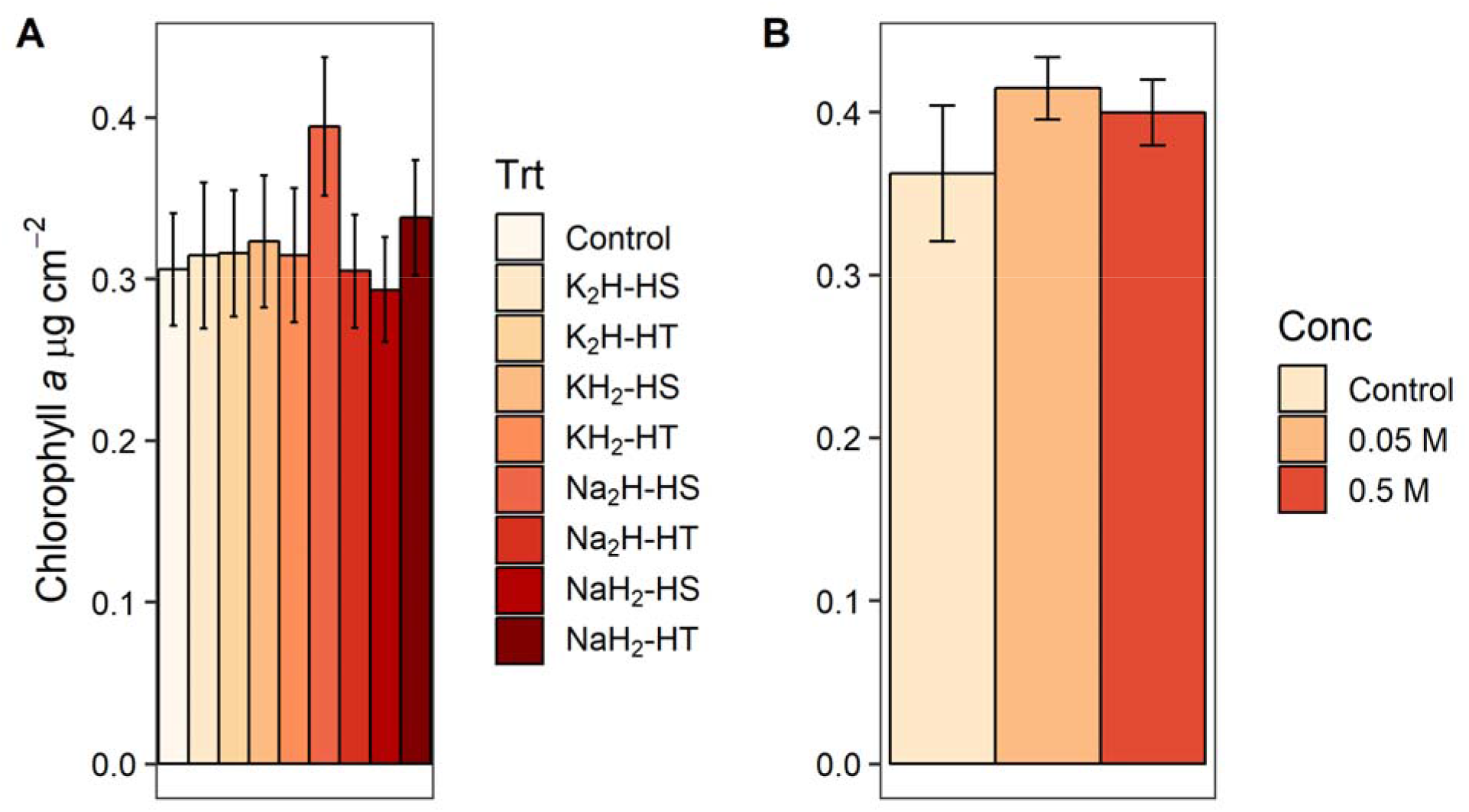
Effect of experimental preparation treatments on chlorophyll *a.* Chlorophyll *a* means ± 1 standard error from NDS field experiments (see Figure 2 caption). (A) For experiments 1-2, total n=107. (B) For experiment 3, total n=105.

**Fig 6.**
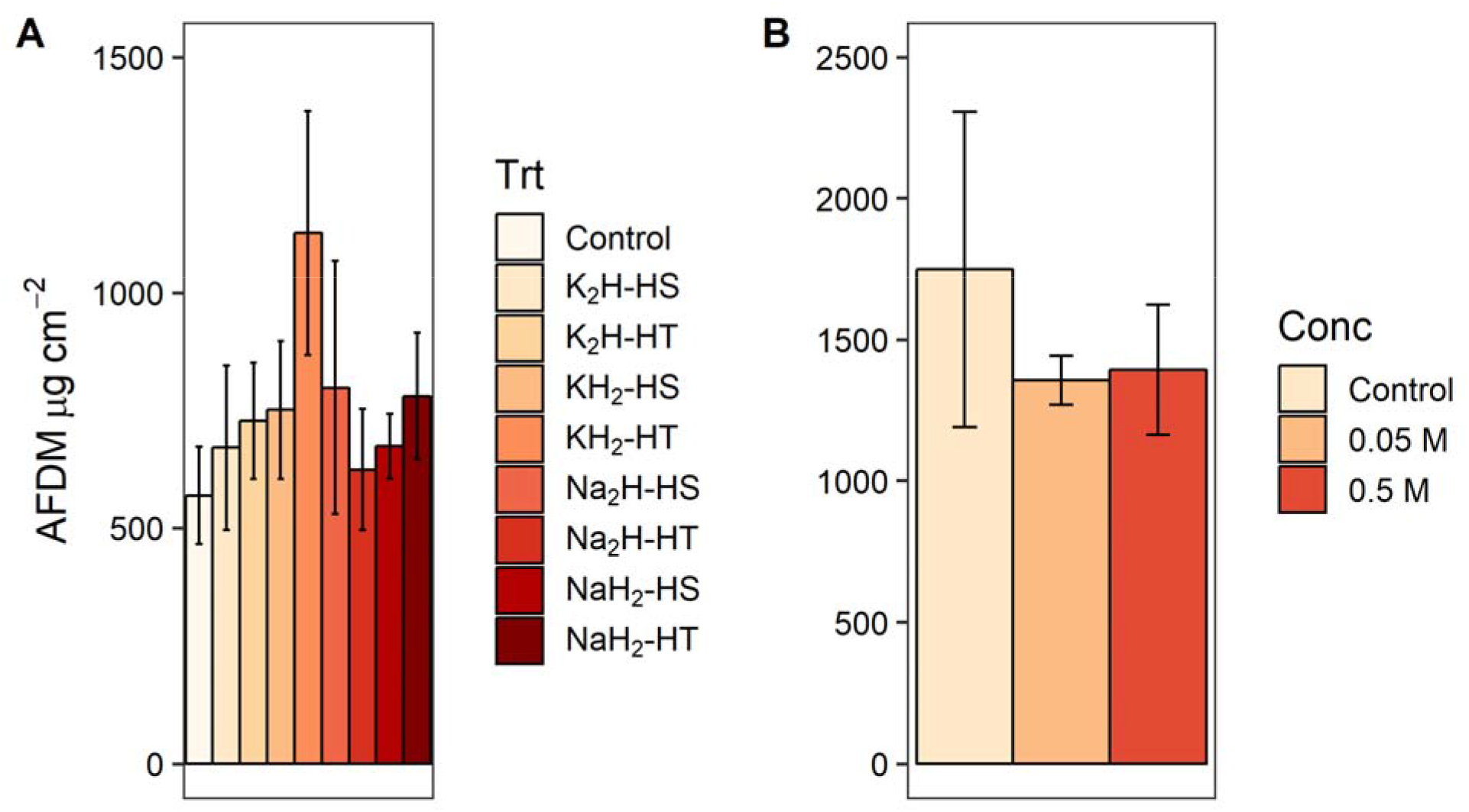
Effect of experimental preparation treatments on ash-free dry mass (AFDM). Ash-free dry mass means ± 1 standard error from NDS field experiments (see Figure 2 caption). (A) For experiments 1-2, total n=93. (B) For experiment 3, total n=104.

**Fig 7.**
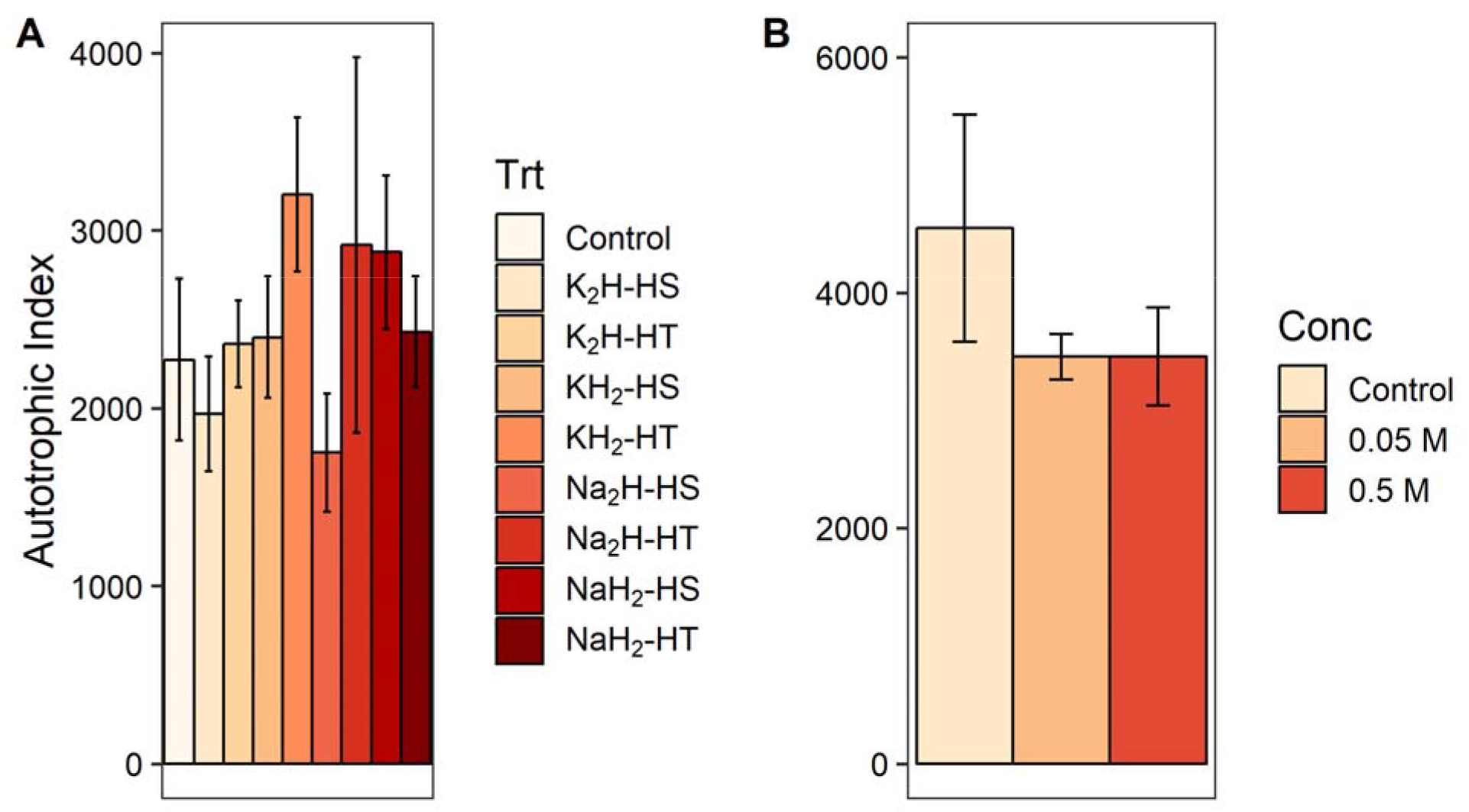
Effect of experimental preparation treatments on an autotrophic index (AI). Autotrophic index means ± 1 standard error from NDS field experiments (see Figure 2 caption). (A) For experiments 1-2, total n=92. (B) For experiment 3, total n=104.

### Experiments

To address the remaining hypothesized mechanisms for inhibition of algal growth by P amendments we conducted a series of field experiments in a small sub-alpine stream. We found no significant differences between any of the eight P preparation treatments (Fig 1) and control treatments (all p >0.05) for any of the algal response variable raw values, indicating neither inhibition nor stimulation of algal growth was induced by the addition of P relative to the control in our experiments.

When we grouped the eight P preparation treatments by three different treatment classes (cation type, phosphate form, and heating method, Fig 1) and tested treatment class effects on response variable effect sizes (S2 Table), we found a significant effect of phosphate form (monobasic or dibasic) on GPP (F_1,88_=5.057, p=0.027) and biomass-specific GPP (F_1,87_=5.578, p=0.020). Dibasic treatments produced higher rates for both measures of primary production (Figs 3-4). Furthermore, phosphate form significantly altered the pH of the agar (F_1,28_=1408.333, p<0.001) and the water (F_1,12_=503.244, p<0.001) in the laboratory experiments. Monobasic chemicals significantly lowered pH means (agar pH=4.81, water pH=5.45) and dibasic chemicals significantly raised pH means (agar pH=8.88, water pH=8.28) relative to controls (agar pH=7.25, water pH=6.35). We did not find an effect of the phosphate cation salt or heating treatment on any algal response variable effect sizes (p>0.05) in the field experiments (Figs 3-7, S2 Table).

We also did not find any significant main effects of phosphate concentration or interactions between concentration and other treatment classes on response variable effect sizes in the field experiments (S3 Table).

## Discussion

Our experiments and quantitative literature analyses did not identify a clear mechanism to explain why 12.9% of past NDS experiments reported a significant negative effect of P treatments on algal growth, a number more than twice as high as the type I error rate (5%) and higher compared to what was observed on N and NP treatments (both <5%). In our experiments we observed slight but non-significant P stimulation rather than inhibition, consistent with the evidence for P-limitation in many freshwater streams [4]. However, past laboratory experiments [17, 18] and a previous meta-analysis [19] support the plausibility of direct P toxicity, cation toxicity, or phosphate form as mechanisms that may inhibit algal growth. Given that multiple mechanisms may be acting simultaneously, and some mechanisms are most likely ecosystem dependent, we discuss the implications of each of our findings and recommend a series of practical steps for future NDS experiments to reduce the likelihood of P-inhibition of algal growth resulting from artifacts of experimental design or experimental preparation.

### P toxicity

While our experiments did not show evidence of P toxicity as a mechanism for P-inhibition of algal growth, previous research supports the possibility of P toxicity occurring across a range of stream ecosystems. In contrast with previous research [7], in our experiments we saw no significant difference in algal biomass between the low (0.05 M) P concentrations and high (0.5 M) P concentrations. Phosphorus toxicity is dependent upon biological and environmental context (see below), and the P concentrations we used in this experiment may have been too low to induce toxicity. There is evidence that NDS experiments in other systems may commonly exceed the concentrations required to detect toxicity, as a previous meta-analysis of 534 NDS experiments showed that higher P concentrations in NDS treatments significantly decreased P effect sizes [19]. The physiological mechanisms that cause P toxicity in algae are not well defined, but some insight can be gained from the terrestrial plant literature.

Terrestrial plant studies have demonstrated that excess P within a cell can induce Fe [15], K [15], or Zn [16] deficiencies. This leads to leaf necrosis and discoloration [33], reduced growth rates [15], and plant death [34, 35]. These plant symptoms have occurred even when studies maintained optimal levels of other nutrients and pH [15], and when multiple phosphate forms have been tested in the same study [36]. Algae [37], like terrestrial plants [38], can take up excess P for storage (i.e. “luxury P uptake”), a strategy to deal with heterogenous nutrient supplies common in stream ecosystems. The mechanisms and potential consequences of luxury P consumption for algae are less clear, but it is plausible that P could accumulate to toxic levels within cells and induce other elemental deficiencies as has been shown for terrestrial plants. However, toxicity from luxury P consumption would only occur if NDS nutrients diffuse at high enough rates that excess P could accumulate in the water-cell boundary layer or biofilm.

Previous studies have measured NDS nutrient diffusion rates in beakers of distilled water to estimate stream water diffusion rates. For instance, a study showed that plastic vial NDS (5.1 cm^2^ area) constructed with 50 mM KH_2_PO_4_ can release 0.321 mM PO_4_^3-^-P·hr^-1^ at day 0, but that diffusion rate declines to 0.001 mM PO_4_^3-^-P·hr^-1^ by day 14 [39]. Clay pot NDS (86.8 cm^2^ area) constructed with 50 mM KH_2_PO_4_ can release 0.113 mM PO_4_^3-^-P ·hr^-1^ P at day 0, with a diffusion rate that declines to 0.011 mM PO_4_^3-^-P ·hr^-1^ by day 14 [39]. Because diffusion rates decline in a log-linear fashion over time, algal populations may initially experience concentrations of P from NDS that are sufficiently high to induce toxicity and inhibition of growth (e.g., 0.019 mM PO_4_^3—^ P [18]). To optimize NDS experiments, future studies could complete pilot experiments that empirically measure diffusion rates to determine appropriate P starting concentrations and experimental lengths (described in detail by Costello et al. [40]). Ultimately, studies should be long enough to surpass the initial high pulses of P released from NDS when direct toxicity may occur, but short enough to maintain a measurable nutrient flux that is significantly enriched from that of the control treatment.

There is also evidence that the availability of other resources can influence P toxicity. Nitrogen- or light-limitation could potentially induce toxicity at lower concentrations of P, if luxury P [37] accumulates without being used for growth due to limitation by other resources. Terrestrial plant studies have shown a positive relationship between P toxicity concentrations and N:P and K:P resource ratios [41], likely because high growth rates supported by N and K availability can reduce tissue P concentrations [36]. Furthermore, P-inhibition of algal growth tends to be stronger in shaded areas [19] where light may be limiting and where NO_3_^-^ reduction could be limited by the availability of NADPH from photosynthesis [42]. Thus, thresholds for P toxicity appear to be closely linked with the probability of secondary limitation by another resource, which would in part explain the inconsistency of reported P-inhibition in NDS studies as environmental conditions change among different ecosystems.

### Cation and Phosphate Form

Few NDS studies have controlled for the effects of phosphate cation or phosphate form (monobasic vs. dibasic). However, experimental evidence suggests that toxicity thresholds differ based on the cation in the phosphate salt [18]. In general, it appears K leads to toxicity at lower phosphate salt concentrations than Na. Growth of the freshwater chrysophyte *Dinobryon sociale* was maintained in the laboratory at P concentrations of 0.032 mM PO_4_^3-^-P for NaH_2_PO_4_ but declined when P concentrations were raised from 0.005 mM PO_4_^3-^-P to 0.019 mM PO_4_^3-^-P for KH_2_PO_4_ [18]. Similarly, a laboratory study on a cyanobacteria, *Microcystis spp.,* showed lower toxicity thresholds for KCl as compared to NaCl [43], further supporting the potential for cation toxicity with the same cation but for different salts. However, across hundreds of published field studies, P effect sizes for algal biomass were higher for K phosphates as compared to Na phosphates [19]. Taken together these results suggest that K-toxicity of algal biomass can be induced under laboratory conditions, but K concentrations in NDS phosphate salts may not be high enough to induce toxicity in field experiments.

Although the cation in the phosphate salt does not appear to strongly influence the effect of NDS P treatments on algal growth, it is likely that the phosphate form (monobasic vs. dibasic) used influences experimental outcomes by modifying pH at the surface of the NDS. In our experimental stream, the pH varied between 7.79 and 8.10, which differed substantially from the agar amended with monobasic phosphates (pH 4.81 ± 0.09) but was more comparable to the agar amended with dibasic phosphates (pH 8.88 ± 0.06). These differences in pH between the P treatments may have contributed to the difference in GPP and biomass-specific GPP we saw between these treatments, i.e., increased productivity on dibasic treatments as compared to the monobasic treatments. These experimental results are consistent with a previous meta-analysis which showed P and NP effect sizes for algal biomass were significantly higher on dibasic NDS treatments relative to those effects on monobasic treatments [19]. Our meta-analysis in this study showed that P-inhibition was reported more commonly than N-inhibition of algal growth, which could be driven by P compounds containing easily disassociated H+ ions while N compounds often do not. To avoid the artifact of pH on NDS P treatments we recommend that NDS experiments mix monobasic and dibasic phosphates to reflect the background pH of study streams as best as possible. This simple step would avoid the confounding influence of alteration of pH in the P treatment of NDS experiments.

### Heating Method

We also investigated the potential for H_2_O_2_ production during the preparation of NDS P treatments to create an artifact in NDS experimental results, as has been hypothesized [44]. We found that heating phosphate and agar together vs. separately on a hotplate did not produce significant differences in algal response metrics. We did not directly test whether H_2_O_2_ was produced in our experiments as was found by a laboratory study that autoclaved phosphate and agar together [21], but either H_2_O_2_ production requires the combination of heat and pressure (from autoclaves) and was not produced in this experiment, or the concentrations of H_2_O_2_ in our experiments were not high enough to inhibit algal growth. A simple solution to avoid the potential inhibiting effect of H_2_O_2_ would be to heat P and agar separately (described by Tank et al. [9]) to avoid the possibility of H_2_O_2_ interference with algal growth. This approach would require little extra effort in NDS experiment preparation and completely remove this potentially confounding artifact.

### Microbial Competition

Indirect mechanisms have also been proposed to explain why P additions commonly inhibit algal growth. Heterotrophic and autotrophic microbial communities interact in complex ways that may change along nutrient gradients. At low concentrations of P, heterotrophic microbes are expected to be competitively dominant because of their strong affinity for P and high surface area relative to volume [45]. However, heterotrophs may also regenerate nutrients that can stimulate autotrophic production and autotrophs produce organic C that fuels heterotrophy, leading to a coupling of the two communities [46, 47]. When nutrients are added to the system these community dynamics are altered, affecting the biomass and diversity of both heterotrophs and autotrophs within the biofilm [48]. A study of Idaho streams found a strongly stimulated AI (i.e., a higher proportion of heterotrophs) when C was added, and a weakly stimulated AI when P was added [22]. Furthermore, a study of Texas streams showed a decoupling of autotrophic and heterotrophic production when nutrients were added [46]. Our experiments and meta-analysis produced no evidence that heterotrophic-autotrophic interactions were influenced by P additions (i.e., no change in AI), but future studies should consider alternative response metrics for measuring heterotrophic microbial biomass if heterotrophic estimates are required to answer study-specific research questions.

### Grazer Selection

In addition to heterotrophic microbes affecting algal growth through ecological interactions, we hypothesized that P effect sizes on grazer exclusion treatments may be larger than on grazed treatments because some grazers have been shown to selectively consume P-enriched resources [25]. Our meta-analysis did not produce evidence to suggest that P effect sizes differed between grazed and ungrazed plots. However, because of the difficulty of conducting grazer exclusion NDS studies, few studies to date have investigated resource selectivity in grazers under field conditions. We recommend that future experiments consider the interactions between nutrient-specific foraging and algal nutrient limitation to determine whether grazing leads to apparent P-inhibition of algal growth. One option is to construct electrical exclusion treatments which can prevent grazing from macroinvertebrates and vertebrates across a wide range of body sizes [49–51].

### Additional Considerations

There are several potential reasons why our experiments did not show significant P treatment effects that have previously been described in the literature. First, deploying only six replicates per treatment produced low statistical power, given the large number of treatments and the small effect sizes of P additions in our NDS experiment. Furthermore, phosphate and cation inhibition of algae have been demonstrated using controlled laboratory experiments that can achieve a wider and more precise concentration gradient as compared to field experiments. We previously observed P-inhibition of algal biomass at the same stream reach used in this study. However, NDS diffusion rates [52] and environmental characteristics in the field are clearly variable over space and time, which may have obscured our ability to connect field-scale results with proposed mechanisms that are largely based on laboratory studies and meta-analyses of field studies.

## Conclusion

Phosphorus additions have significantly inhibited algal biomass in 12.9% of past NDS studies, and investigators have hypothesized that this may be an artifact of NDS preparation or an indirect effect of increased heterotrophic microbial competition or top-down grazer control. We found that phosphate form (monobasic vs. dibasic) likely influences algal growth on NDS P treatments by mediating biofilm pH levels, and acidic monobasic treatments may inhibit algal growth. Furthermore, the literature supports direct P toxicity as a mechanism for P-inhibition of algal growth [17–19], particularly when other resources such as light or N are limiting [37]. We did not find support for phosphate salt cation toxicity occurring under field conditions, nor did we find evidence that laboratory heating method influenced algal responses to P. Based on our analyses, it is also unlikely that P stimulates heterotrophic microbes relative to autotrophic microbes or that P stimulates grazing rates.

Considering that multiple mechanisms may be operating simultaneously to inhibit algal growth on NDS P treatments, we recommend several low effort, cost-effective steps for the NDS preparation process that could reduce the potential for P-inhibition in future experiments. First, future experiments could measure background stream nutrient concentrations to determine the most appropriate P treatment concentrations for NDS construction [53], with the goal of avoiding the potential for levels of P that are directly toxic to algae. In addition, measuring NDS diffusion rates [39, 40, 54] under conditions that mimic natural systems would allow investigators to further determine appropriate concentrations and experimental lengths for NDS studies. Experiments should be long enough to surpass the potential for initial P toxicity when diffusion concentrations are at their highest but short enough to maintain stimulatory P diffusion from NDS. Because monobasic and dibasic phosphates influence NDS pH levels, we encourage investigators to mix the two phosphate forms to reflect the background pH in experimental streams to the extent possible. Finally, while we did not find evidence that laboratory construction methods inhibited algal growth, it seems prudent (and logistically simple) to use the separate agar and phosphate heating methods outlined by Tank et al. [9] to avoid the potential for H_2_O_2_ production that inhibits microbial growth [21]. Avoiding confounding factors in NDS experiments will ensure that studies are not underestimating P-limitation of primary producers in aquatic ecosystems, improving our understanding of how resources and environmental conditions interact to affect algal growth and stream ecosystems as a whole.

## Acknowledgements

This manuscript benefited from reviews by Keeley MacNeill and Steven Francoeur. We also thank Mitchell Ralson for help in the field and laboratory.

## Supporting Information Captions

**S1 Table. Environmental variables measured during nutrient diffusing substrate experiments at Little Beaver Creek, CO.** Multiple measurements are recorded as mean ± standard deviation. Several nutrient measurements were below detection. pH was not measured during experiment 3 because of an instrument malfunction.

**S2 Table. Results of ANOVAs testing the effect of three treatment classes on P effect sizes.**

Four different phosphate chemicals (KH_2_PO_4_, K_2_HPO_4_, NaH_2_PO_4_, Na_2_HPO_4_) crossed with laboratory heating methodology were deployed in two NDS experiments in 2016. Treatment classes in the statistical analyses included phosphate cation, phosphate form, and heating method. ANOVA results testing the effect of each treatment class on each response variable’s P treatment effect size are presented, with experiment included as a fixed effect block in all models. P<0.05 is indicated as “*”.

**S3 Table. Results of ANOVAs testing the effect of four treatment classes on P effect sizes.**

Four different phosphate chemicals (KH_2_PO_4_, K_2_HPO_4_, NaH_2_PO_4_, Na_2_HPO_4_) crossed with laboratory heating methodology and two phosphate concentrations were deployed in an NDS experiment in 2017. Treatment classes in the statistical analyses included phosphate cation, phosphate form, and heating method, along with concentration as a potential modifier. Two-way ANOVA results are presented where each treatment class was crossed with concentration, to determine effects on each response variable’s P treatment effect size. P<0.05 is indicated as “*”.

**S1 References. Studies used in autotrophic index meta-analysis.**

**S2 References. Studies used in grazer exclusion meta-analysis.**

